# Cellular rejuvenation protects neurons from inflammation mediated cell death

**DOI:** 10.1101/2023.09.30.560301

**Authors:** Sienna S. Drake, Abdulshakour Mohammadnia, Kali Heale, Adam M.R. Groh, Elizabeth M.-L. Hua, Aliyah Zaman, Matthew A. Hintermayer, Stephanie Zandee, David Gosselin, Jo Anne Stratton, David A. Sinclair, Alyson E. Fournier

## Abstract

In multiple sclerosis (MS), the invasion of the central nervous system by peripheral immune cells is followed by the activation of resident microglia and astrocytes. This cascade of events results in demyelination, which triggers neuronal damage and death. The molecular signals in neurons responsible for this damage are not yet fully characterized. In MS, retinal ganglion cell neurons (RGCs) of the central nervous system (CNS) undergo axonal injury and cell death. This phenomenon is mirrored in the experimental autoimmune encephalomyelitis (EAE) mouse model of MS. To understand the molecular landscape, we isolated RGCs from mice subjected to the EAE protocol. RNA-sequencing and ATAC-sequencing analyses were performed. Pathway analysis of the RNA-sequencing data revealed that RGCs displayed a molecular signature, similar to aged neurons, showcasing features of senescence. Single-nucleus RNA-sequencing analysis of neurons from human MS patients revealed a comparable senescence-like phenotype., which was supported by immunostaining RGCs in EAE mice. These changes include alterations to the nuclear envelope, modifications in chromatin marks, and accumulation of DNA damage. Transduction of RGCs with an *Oct4*-*Sox2*-*Klf4* transgene to convert neurons in the EAE model to a more youthful epigenetic and transcriptomic state enhanced the survival of RGCs. Collectively, this research uncovers a previously unidentified senescent-like phenotype in neurons under pathological inflammation and neurons from MS patients. The rejuvenation of this aged transcriptome improved visual acuity and neuronal survival in the EAE model supporting the idea that age rejuvenation therapies and senotherapeutic agents could offer a direct means of neuroprotection in autoimmune disorders.

## Introduction

MS is an immune mediated disease with a neurodegenerative component affecting approximately 2.3 million people worldwide^1^. While immune-mediated demyelination of axons constitutes a primary pathogenic mechanism in MS, sustained clinical deficits are associated with neuronal degeneration including loss of neurons in the gray matter and loss of axons in white matter lesions and in normal appearing white matter^2-4^. Analysis of both non-lesioned and lesioned post-mortem brain, spinal cord and optic nerve have provided evidence of extensive and diffuse axonal degeneration present throughout the CNS of MS patients^5-12^. Axonal pathology appears early on in disease, suggesting neurodegeneration may occur in parallel with demyelination. Multiple complex contributors seem to drive axonal and neuronal degeneration in MS. For example, loss of myelin sheath trophic support, noxious factors released by activated glia, infiltrating immune cells and degenerating myelin, iron neurotoxicity, mitochondrial dysfunction and oxidative stress induced by reactive oxygen and reactive nitrogen species, and glutamate excitotoxicity all are evidenced to contribute to axonal and neuronal cell body injury in MS^13-17^.

It is certain that the complex inflamed, and injurious environment alters intrinsic neuronal gene programs, as it does to the CNS glial cells, and these altered gene programs may contribute to or protect from neurodegeneration in the disease^18^. Animal models of disease are useful for assaying neuronal changes in response to pathological inflammation, enabling sampling from various timepoints across a standardized disease course. One model is the experimental autoimmune encephalomyelitis (EAE) mouse which recapitulates multiple aspects of MS neuronal pathology, particularly in the retina and optic nerve. Here, EAE mice exhibit demyelinating lesions of the optic nerve, immune cell infiltration and gliosis, retinal nerve layer thinning and neuronal death, axonal pathology, and loss of visual acuity throughout the disease course, features which are likewise observed in MS patients^19^. It stands to reason that conducting RNA-sequencing of retinal ganglion cells (RGCs) from EAE mice and comparing these data with preexisting sequencing datasets of neurons from MS patients may provide insight into a conserved neuronal response to pathological inflammation.

## Results

### The RGC transcriptome in EAE resembles naïve aged RGCs

To better understand the response of neurons to pathological inflammation, retinal ganglion cells (RGCs) were collected by fluorescence-activated cell sorting (FACS) from EAE-(2-3 months old), naïve age-matched-(2-3 months old), and naïve aged-(6-7 months old) Thy1-vGlut2-YFP transgenic mice. Sorted RGC populations were sequenced and DESeq2 was used to obtain normalized read counts and compare gene fold changes between EAE samples and naïve age-matched samples or naïve aged samples. Linear regression analysis of the log2FC values identified a significant correlation between the fold change of all detected genes in RGCs between EAE and naïve-aged mice suggesting that EAE neurons exhibit an aged signature (R^2^=0.47, pval < 0.001, Fig. 1A). Assessing all significantly dysregulated genes, over 50% of the upregulated genes in EAE mice were also significantly upregulated in naïve 6-month-old mice and over 50% of genes significantly downregulated in EAE RGCs were also downregulated in aged RGCs further supporting a common signature of gene expression patterns between young EAE and old naïve RGCs (Fig. 1B). To validate this finding, an RNA-sequencing dataset comparing 12-month-old RGCs to 5-month-old RGCs was obtained from the NIH BioProject (Accession ID: PRJNA655981), reanalysed following the same workflow to obtain gene expression changes between 12 month-old and 5-month-old RGCs, and a list of the top 100 significantly upregulated genes in the aged RGCs was generated^20^. Here, ranked gene set analysis showed significant enrichment of this gene list in both the 6 month aged RGCs and EAE RGCs, further suggesting that genes upregulated in aging RGCs are likewise being upregulated in young inflamed RGCs (Fig. 1C,D). Cellular aging, termed senescence, is a collection of altered cellular pathways and dysregulated gene expression that often arise in aged cells and has been predominantly characterized in non-neuronal cells where ongoing cell division, causing replicative stress, induces departure from the cell cycle^21^. Although neurons have undergone terminal differentiation, aged neurons exhibit characteristics of senescence including DNA damage and elevated expression of proinflammatory molecules^22,23^. To compare our neuronal signature to classically defined senescence genes, ranked gene set enrichment was conducted comparing the EAE RGCs, six-month RGCs, and aged RGC data to a curated senescence gene set. This gene set (SEN_MAYO) contains genes known to be involved in senescence and commonly upregulated in aging, with classifications as cytokine/chemokines, transmembrane, or intracellular proteins and is consistently enriched in various aged tissue samples across mice and humans^24^. Here, ranked gene set enrichment found significant enrichment of the SEN_MAYO gene set in all three gene expression datasets – EAE, 6-month-old, and 12-month-old RGCs (Fig. 1E-G). Indeed, the 12-month-old RGCs had the largest normalized enrichment score (NES=2.12), followed by EAE RGCs (NES=1.72), and finally by 6-month-old RGCs (NES=1.38), suggesting that inflamed RGCs exhibit features of senescence (Fig. 1E-G). One common feature of senescence is the senescence associated secretory phenotype (SASP), which consists of the upregulation of a group of secreted inflammatory factors. Thus, the SEN_MAYO gene set was subsetted based on the genes’ prior classification as cytokines/chemokines (CYTO) to evaluate the contribution of inflammatory secreted molecules to the senescence phenotype, compared to the contribution of the non-cytokine/chemokine genes (transmembrane and intracellular classification, NOT_CYTO). While there was a significant contribution of cytokines/chemokines to the EAE signature (Fig. 1E), both SEN_MAYO gene subsets were significantly enriched suggesting that both inflammatory genes and other genes are contributing to the senescence-like phenotype in EAE neurons (Fig. 1E). In 12-month-old RGCs, the CYTO genes subset made a comparatively smaller contribution (p=0.05), which affirms the stronger inflammatory component of EAE compared to natural aging (Fig. 1F). In the 6-month-old RGCs, neither subset on its own reached statistical significance, evidencing the lower level of senescence at this age (Fig. 1G). Finally, an RNA-sequencing dataset from motor neurons of EAE mice was evaluated for the SEN_MAYO gene sets (BioProject Accession ID PRJNA414103)^25^. Similarly, the SEN_MAYO gene set was significantly enriched in EAE motor neurons, with significant enrichment from both the CYTO and NOT_CYTO subsets (Fig. 1H). Overall, these data demonstrate that inflamed RGCs are transcriptionally older and acquire a senescence associated gene signature, that is likewise present in EAE motor neurons, and consists of both non-secreted proteins and secreted inflammatory proteins.

**Figure 1.**
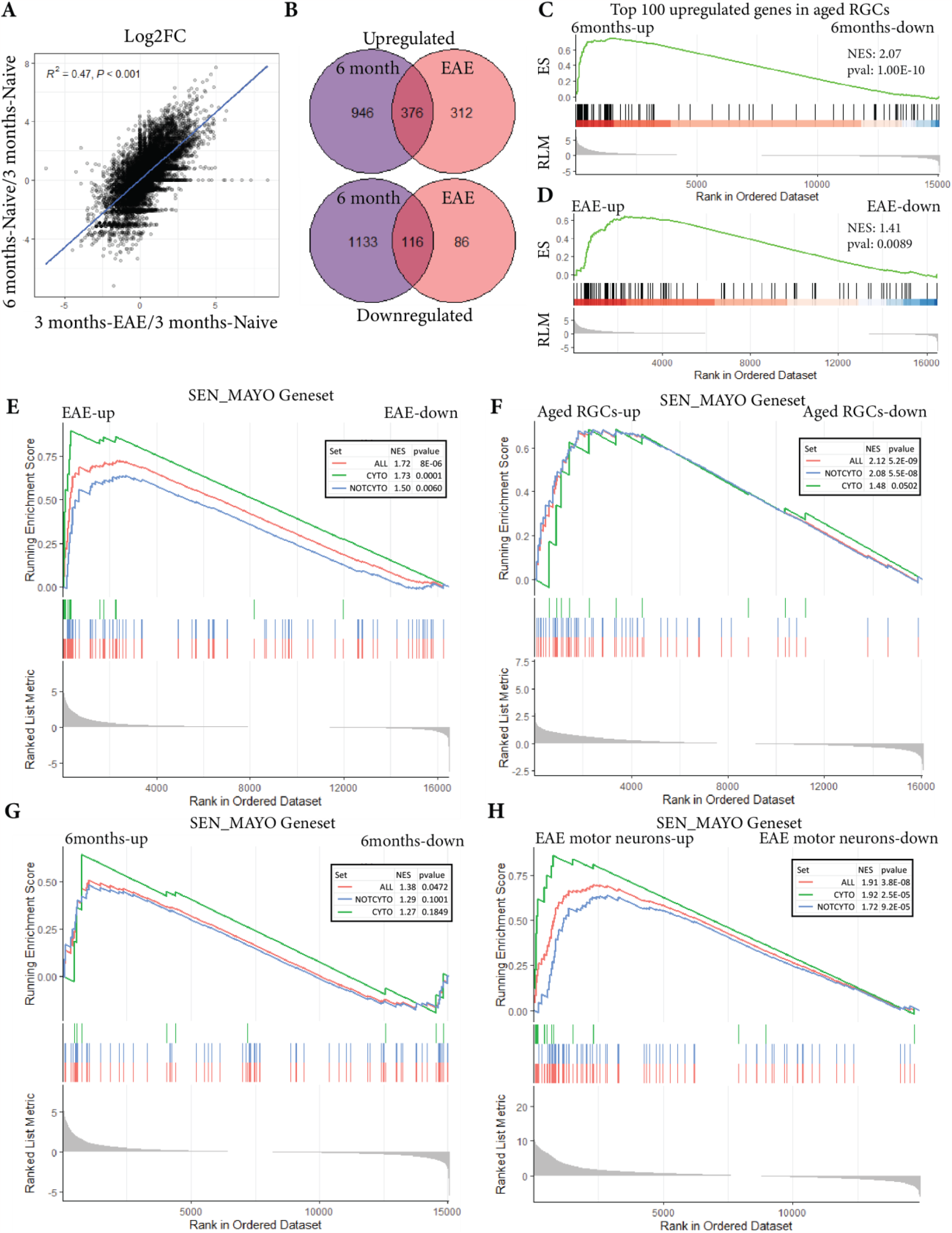
Age-associated changes in transcriptome of RGCs from EAE mice. A) log2FC for each gene of 6 month-old naïve RGCs (y axis) vs 3 month-old EAE RGCs (x axis), with a linear regression curve fit to the data. B) Overlap between significantly (padj < 0.05) upregulated (top) and downregulated (bottom) genes in 6 month-old naïve and 3 month-old EAE RGCs, with the number in the bubbles referring to the count of genes in each set. C-D) Ranked gene set enrichment result of the top 100 upregulated genes in aged RGCs for 6 month old (C) and EAE RGCs (D). E-H) Ranked gene set enrichment result for the SEN_MAYO gene set, split by the classification Cytokine/Chemokine, for EAE (E), 6-months old (F), aged RGCs (G), and EAE motor neuron (H) samples. ES=Enrichment Score. RLM=Ranked List Metric, NES=Normalized Enrichment Score.

### Gene co-expression across the EAE disease course identifies altered pathways relevant to senescence

To more fully describe how gene expression was changing in RGCs during EAE, gene co-expression analysis was conducted comparing 3-month-old naïve, 3-month-old presymptomatic, and 3-month-old symptomatic EAE RGC samples. Eight gene modules with at least 300 gene members each were identified, grouping genes with different patterns of expression across the disease course (Fig. 2A). Using ATAC-sequencing data generated from FACS enriched RGCs, the accessibility of the transcription start site (TSS) of all module genes was subdivided into quartiles ranging from least accessible (quartile 1) to most accessible (quartile 4). The genes present in the eight modules were enriched for highly accessible genes (predominantly quartiles 4 and 3), indicating that these are true RGC-expressed genes based on their expression and accessibility (Fig. 2B). Next, GO Biological Process (BP) term enrichment was conducted for each module. A large portion of enriched GO terms were unique to each module, with some overlap, as expected due to the unique gene sets of each module (Fig. 2C). Based on the hallmarks of aging, GO terms related to these processes were assessed in each module, with a focus on those related to DNA damage, autophagy and proteostasis, mitochondrial energy maintenance, cell cycle, senescence, and histone modifications^26^ (Fig. 2D). DNA damage and cell cycle associated pathways were particularly enriched in Module 4, which consists of genes that increase expression at the symptomatic phase of EAE. Pathways related to histone modifications were enriched in Module 2, which consists of genes that decrease throughout presymptomatic and symptomatic EAE. Module 3, consisting of genes that are increased at both presymptomatic and symptomatic, demonstrated significant enrichment of terms related to cellular senescence, and had a preponderance of pathways relevant to inflammation and chemotaxis. Module 1, with genes that increase at symptomatic EAE, demonstrated an enrichment of mitochondrial and energy associated pathways, as well as pathways related to unfolded protein response, cell stress, and apoptosis. Module 5 and Module 7, which decrease in symptomatic EAE, demonstrated an enrichment of pathways related to neuronal processes such as synaptic transmission and dendrite morphology (Fig. 2D,E). Lastly, Modules 6 and 8 consisted of genes that were up and down at presymptomatic EAE respectively, with Module 6 consisting of terms related to protein folding and localization, and module 8 with terms related to mRNA splicing (Fig. 2E). In sum, gene groups that increased were enriched for terms related to inflammation, cell stress, cell cycle and senescence, and DNA damage response, whereas gene groups that decreased were enriched for pathways related to epigenetic regulation and neurotransmission. Together, these results provide further evidence for alterations in senescence-associated processes in RGCs.

**Figure 2.**
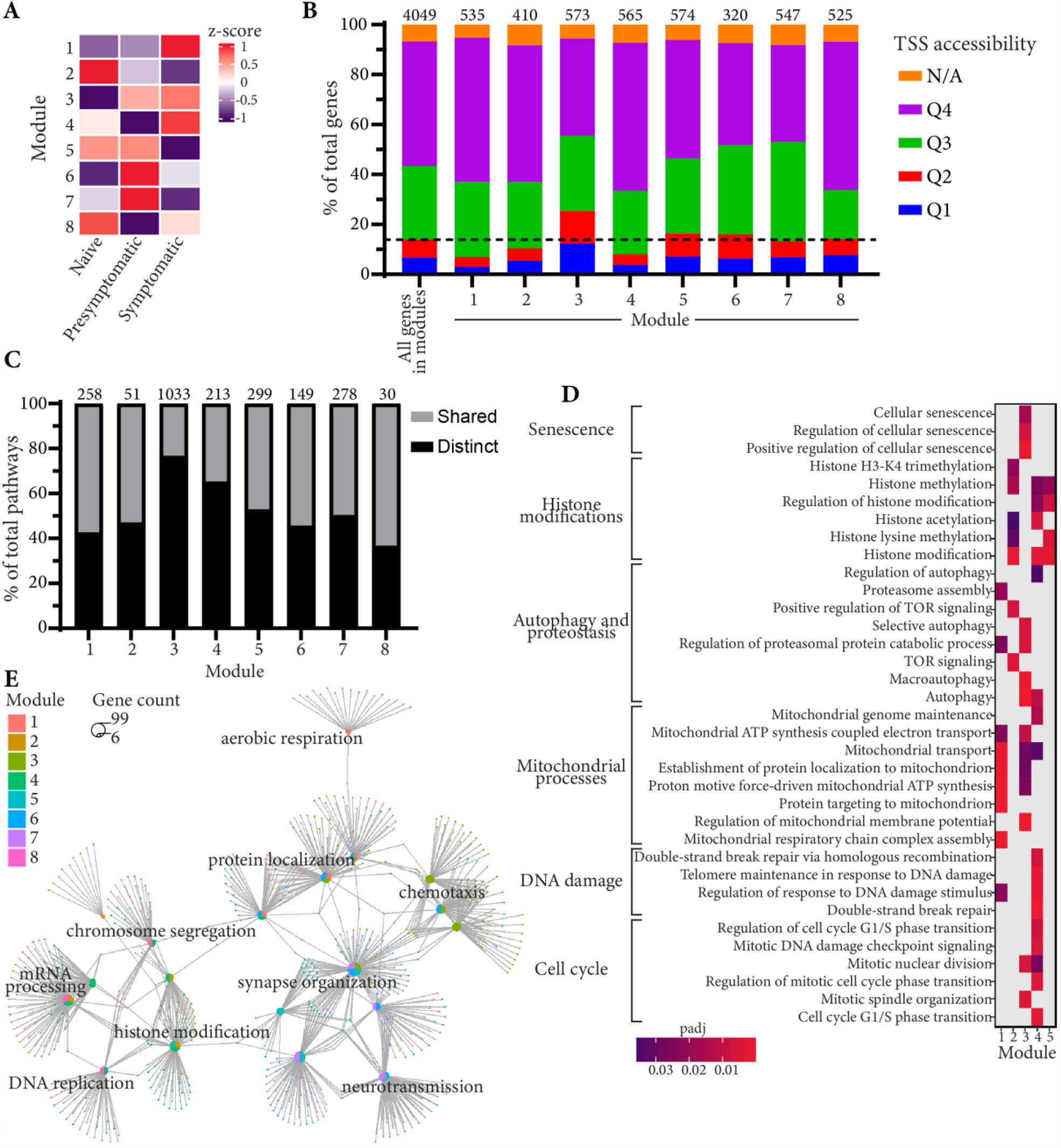
Gene co-expression across EAE disease course identifies altered pathways in EAE. A) Leiden based clustering of genes that change from naïve, presymptomatic, to symptomatic EAE identified 8 gene co-expression modules. B) Transcription start site (TSS) accessibility in quartiles of genes assigned to modules, count of total genes per module listed on top of the bar for each module. C) Percent of distinct significantly enriched (padj < 0.05) GO Biological Processes for each module, count of total number of pathways per module listed at the top of each bar. D) Subset of significantly enriched GO Biological Processes displayed related to pathways involved in senescence, showing enrichment across different modules. E) Simplified network diagram showing top GO terms per module and their relationships to other modules.

### A senescence signature is detected in MS patient neurons

To determine whether the senescent signature detected in the EAE RNA-sequencing samples was likewise enriched in MS, an online available single-nucleus sequencing dataset containing neurons from control and MS patients was accessed and reanalysed to assess average expression across all identified neuronal clusters present in both control and MS patients. Here, 10 neuronal cell clusters were identified, 8 of which had more than 5 cells present in both MS and control samples, which were used for downstream expression averaging and analysis (Fig. 3A). Average neuronal gene expression was computed for all neurons per patient, and then single sample GSEA was conducted to calculate an enrichment score for the SEN_MAYO gene set in each patient, as well as the CYTO and NOT_CYTO subsets. Here, MS patient neurons demonstrated significant enrichment of the SEN_MAYO gene set compared to control patient neurons. Both CYTO and NOT_CYTO subsets were significantly enriched in MS patient neurons revealing a contribution of both non-secreted senescence associated factors and secreted inflammatory senescence associated genes (Fig. 3B). GO term analysis for biological processes related to senescence, DNA damage, cell cycle, and epigenetic modifications were evaluated, and the overlap of de-enriched and enriched pathways in MS patients were compared to the GO terms enriched in the gene modules with decreased expression in symptomatic EAE (Modules 2, 5, and 7) and increased expression in symptomatic EAE (Modules 1, 3, and 4) respectively (Fig. 3C,D). Despite comparing between bulk-sequencing and single-nucleus sequencing, and comparing human to mouse gene sets, there were common pathways identified between the two, particularly for the enriched pathway group where almost a quarter of pathways occurred in at least one of the related EAE modules (Fig. 3D). Notably, pathway overlap was observed between pathways enriched in MS patient neurons and EAE Module 3 (67 pathways, 12% of total), which consisted of immune system and inflammation related GO terms (Fig. 3D). Overlap with EAE Module 1 and Module 4 contributed an additional 4% and 3% of pathways respectively, here with M1/MS overlap consisting of terms like ‘response to unfolded protein’ and ‘ER overload response’ and M4/MS overlap consisting of terms including ‘positive regulation of DNA repair’ and positive regulation of telomere maintenance in response to DNA damage’. 16 pathways (3%) were common to Module 1, Module 3, and the MS neurons including terms including ‘cellular respiration’ and ‘ATP biosynthetic process’. Additionally, numerous pathways were significantly dysregulated related to the same hallmarks of aging: DNA damage, cell cycle and senescence, mitochondrial dysfunction, and histone modifications (Fig. 3E-H). This suggests that, similar to EAE, neurons in MS demonstrate an enrichment of genes related to senescence, and alteration of pathways relevant to the process including DNA damage, epigenetic modifications, and mitochondrial dysfunction.

**Figure 3.**
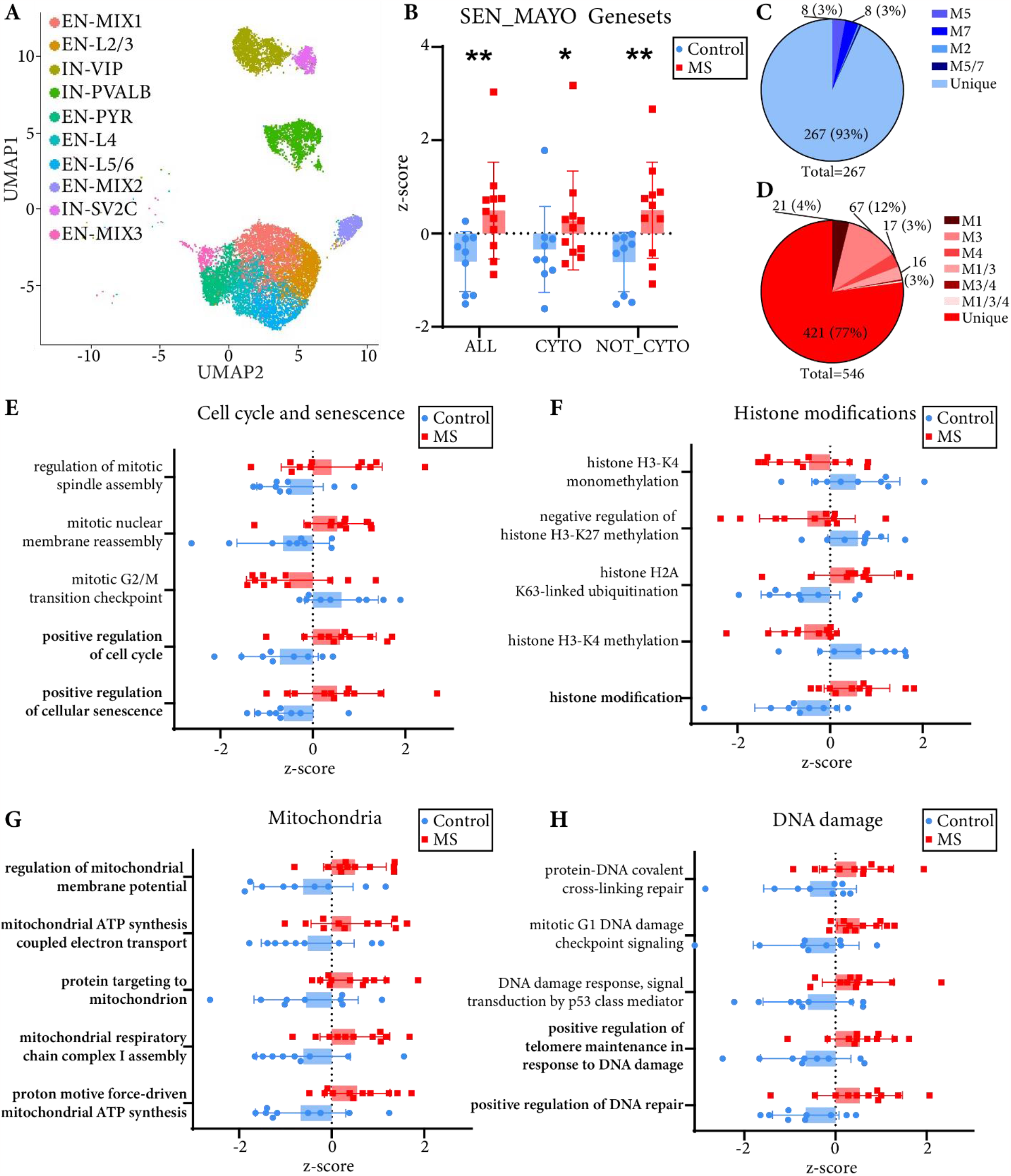
Single-nucleus sequencing of neurons from MS patients identifies senescence associated changes. A) UMAP representation of ten neuronal cell clusters identified from single-nucleus sequencing of MS and control gray matter in Schirmer et al 2019. Clusters EN-MIX2 and EN-MIX3 were only identified in MS patients (5 or less cells in control patient samples) and were excluded from the downstream analysis. B) SEN_MAYO gene set enrichment in MS patients compared to controls. * < 0.05, ** < 0.01, one-tailed Mann-Whitney test. C) De-enriched GO BP terms in MS patients compared to controls and their overlap with EAE Module 2 (M2), Module 5 (M5), and Module 7 (M7), comparisons with overlap of >= 3% are annotated. D) Enriched GO BP terms in MS patients compared to controls and their overlap with EAE Module 1 (M1), Module 3 (M3), and Module 4 (M4), comparisons with overlap of >= 3% are annotated. E-H) Subset of significantly dysregulated GO terms related to cell cycle and senescence (E), histone modifications (F), mitochondria (G), and DNA damage (H), all significant at padj < 0.05. Bolded terms reflect their significant enrichment in one or more EAE modules.

### IHC of EAE retinal cross-sections identified dysregulated protein signatures relevant to senescence

With a transcriptomic signature suggesting senescence-related processes in EAE neurons, protein markers of senescence were assessed by IHC in EAE retinal cross-sections to validate that these pathway level changes were reflected at a protein level. One common feature of senescence is the accumulation of DNA damage, which likewise occurs in aged neurons^23,27^. Given the evidence for alterations to DNA damage pathways at the transcriptomic level, this was assessed in EAE retinal tissue through yH2AX immunoreactivity (Fig. A,B). Here, there was a significant increase in the number of BRN3A positive RGC nuclei with γH2AX positivity in EAE RGCs compared to naïve age matched mice (Fig. 4C). Next, histone modifications were evaluated by IHC for H3K27 trimethylation (Fig. 4D,E). While this mark may be up or downregulated in senescent cells, there is evidence that in aged neurons it is upregulated^28^. Similarly, increased fluorescence intensity of nuclear H3K27me3 was observed in the EAE animals, notably at peak disease, compared to control mice (Fig. 4F). Another common feature of senescent cells is the loss of nuclear envelope integrity, evidenced by IHC for nuclear lamina proteins^29,30^. Here, LAMINB1 IHC was conducted to assess the nuclear lamina morphology in RGCs in EAE (Fig. 4G,H). Compared to naïve control mice, EAE mice had an increased number of RGCs with abnormal or dystrophic nuclear envelopes (Fig. 4I). Together, these data suggest that RGCs in EAE mice demonstrate dysregulation of proteins related to aging and senescence.

**Figure 4.**
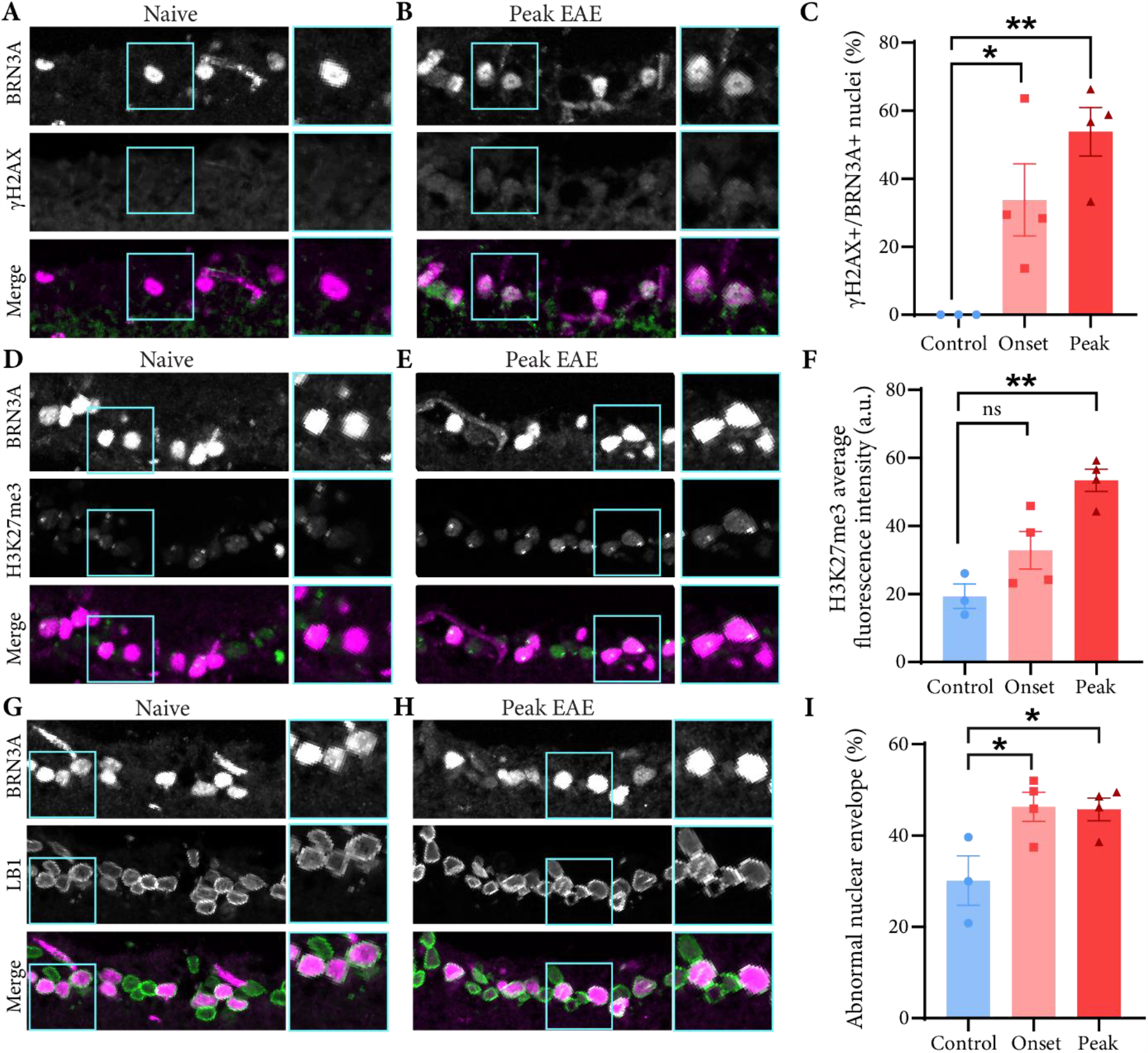
Immunohistochemical analysis of EAE retina detects changes in senescence associated protein markers. A-C) Immunohistochemical analysis of γH2AX in the RGC layer of naïve (A) and peak EAE (B) mice. BRN3A in magenta labels RGCs, and γH2AX in green (A,B) is detected in the nuclei of RGCs in EAE mice (C). D-F) Immunohistochemical analysis of H3K27me3 in the RGC layer of naïve (D) and peak EAE (E) mice. BRN3A in magenta labels RGCs, and H3K27me3 in green (D,E), with average nuclear fluorescence intensity quantified in (F). G-I) Immunohistochemical analysis of LAMINB1 (LB1) in the RGC layer of naïve (G) and peak EAE (H) mice. Abnormal nuclear envelopes, as defined by incomplete or invaginated rings is quantified in (I). * < 0.05, ** < 0.01, unpaired student’s t-test.

### Rejuvenation of RGCs by expression of *Oct4, Sox2*, and *Klf4* prevents RGC death in EAE

Given the remarkable age-related and senescent signature detected by RNA-sequencing and IHC in the RGCs of EAE mice, an AAV-gene-therapy approach was employed to rejuvenate RGCs to assess whether combating this senescent-like phenotype would aid or harm neurons in response to inflammatory injury in this model. This approach was previously shown to restore a youthful epigenetic and transcriptomic signature to aged RGCs and protect from cell death in an age-related neurodegenerative disease model of glaucoma^20^. AAV2-GFP and AAV2-OSK (vector containing an *Oct4*-*Sox2*-*Klf4* transgene), serotyped to target RGCs, were injected intravitreally 3 days prior to EAE induction (days post induction; dpi). Animals were monitored daily for motor symptom scoring from 7 dpi until 18 dpi at which time the tissue was collected for viral transduction assessment in the retina and RGC survival quantification (Fig. 5A,B). No difference was observed in EAE motor score progression or severity, as expected due to the virus being specific to the RGCs, and suggestive that there are no systemic off target effects on the immune system (Fig. 5B). However, visual acuity, assessed by optomotor response, was significantly higher in the AAV2-OSK compared to AAV2-GFP mice at peak disease. Next, retinal flat-mounts were evaluated by IHC for GFP to label AAV2-GFP transduced cells, or by IHC for KLF4 to label AAV2-OSK transduced cells (Fig. 5C). Then, GFP+ and KLF4+ RGCs were quantified by colocalization of the GFP or KLF4 signal with the RGC protein marker RBPMS. Compared to naïve AAV2-GFP animals, EAE AAV2-GFP animals demonstrated a significant reduction in the number of GFP+ RGCs, suggesting that these RGCs are dying throughout the disease course, comparable to previous reports of RGC cell loss in EAE^31^. In contrast, EAE AAV2-OSK retinas retained the same number of transduced cells as their naïve counterparts, suggesting that AAV2-OSK transduced RGCs survive longer in EAE. Indeed, overall quantification of the total number of RGCs across the retinae found that EAE AAV2-OSK retinas had significantly more RGCs compared with EAE AAV2-GFP retinae. Together, these data suggest that employing an intervention that promotes youthful gene expression maintains function of and protects RGCs during inflammation mediated cell death.

**Figure 5.**
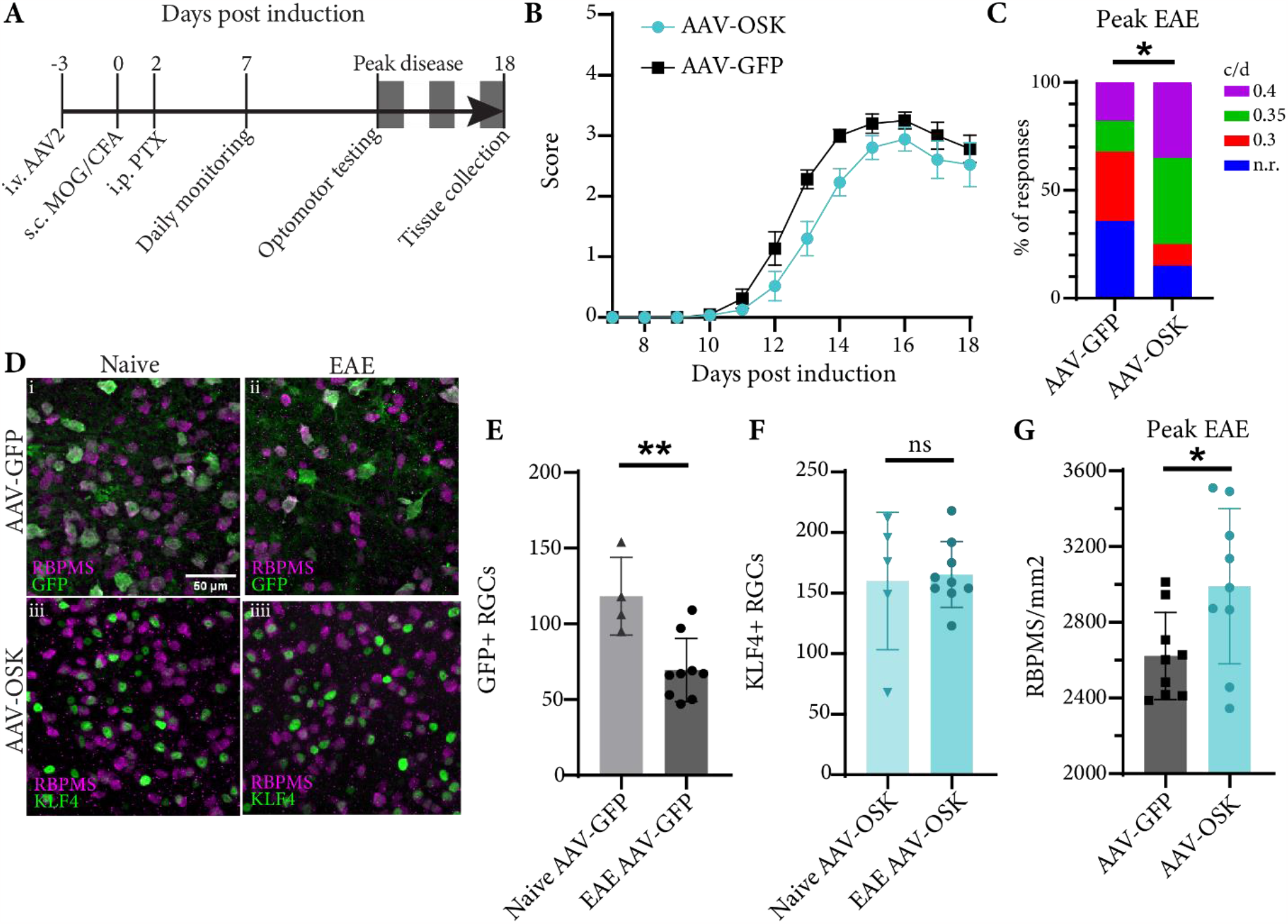
AAV-OSK mediated rejuvenation of RGCs protects from cell death in EAE. A) Diagram of experimental timeline showing AAV-GFP/AAV-OSK bilateral intravitreal injection (i.v.) 3 days pre-EAE induction (d.p.i.), followed by EAE induction through MOG/CFA subcutaneous (s.c.) injection and pertussis toxin intraperitoneal (PTX, i.p.) injection, and daily monitoring of EAE score beginning at 7 dpi and continuing until experiment end at 18 dpi. B) EAE disease course for animals receiving AAV-GFP and AAV-OSK, demonstrating no difference in motor score severity. C) Visual acuity testing by optomotor response quantification demonstrates significant difference between AAV-GFP and AAV-OSK mice at peak EAE disease. c/d = cycles/degree, n.r.=no response at tested levels. * < 0.05, Chi-square test. AAV-GFP n = 28, AAV-OSK n=20. D) Representative images of retinal flatmounts of AAV-GFP (i/ii) and AAV-OSK (iii/iiii) naïve and EAE mice showing RBPMS labeling retinal ganglion cells, GFP labeling virus transduced GFP RGCs, and KLF4 labeling OSK virus transduced RGCs. E) Count of GFP+/RBPMS+ cells in highest density region of retinal flatmount, demonstrating a reduction in GFP+ RGCs in EAE compared to naïve mice. F) Count of KLF4+/RBPMS+ cells in highest density region of retinal flatmount, demonstrating no change in the number of OSK transduced RGCs between naïve and EAE mice. G) Count of total RBPMS+ cells quantified across the retina in EAE AAV-OSK and EAE AAV-GFP control mice, demonstrating greater number of RGCs remaining in retina of EAE AAV-OSK receiving mice compared to control. * < 0.05, ** < 0.01, unpaired student’s t-test.

## Discussion

While aging and senescence are not the same, it is known that aging and age-related injury can induce senescence in cells and that depleting senescent cells can improve age-related diseases, thus they are closely intertwined^21,32^. The phenotype of senescent cells is varied and complex including features such as DNA damage foci, nuclear lamina dysfunction, epigenetic modifications, and a SASP^21^. However, due in part to its initial discovery as an adaptation of proliferative cells to stress and stop dividing, it is less clear what a senescent neuron might look like given its post-mitotic nature^21^. Here we took a dual approach, acquiring evidence by comparing to aging neurons and assessing gene sets and phenotypes related to senescence, as it is expected that a senescent neuronal phenotype would be more common in neurons from aged animals. The finding that inflamed neurons from EAE mice both appear transcriptionally alike to neurons from naïve aged mice, and that both EAE and aged RGCs exhibit an enrichment for senescence genes and processes suggest that inflammation mediated injury to the central nervous system can promote a premature aging and senescent-like phenotype in neurons. To our knowledge, this is the first study demonstrating that CNS autoimmunity may drive neuronal senescence. However, the exact mediators and components of the MS and EAE inflammatory environment that drive neuronal senescence and through what pathways remains to be clarified. Importantly, this neuronal phenotype consists of an inflammatory profile of cytokine and chemokine upregulation which may promote a positive feedback cycle between the acute inflammatory injury in MS lesions and recruitment, engagement, and maintenance of inflammatory mediators throughout the CNS. Notably, this signature is not specific to any single neuronal cell subtype but seems to affect diverse neuronal cell types including RGCs and motor neurons in EAE, and both excitatory and inhibitory cortical neurons in MS patients.

Indeed, the systematic evaluation of age-related changes and senescence in neurons is a relatively new endeavor, however evidence exists for a senescent phenotype^33^. Neurons aged both *in vitro* and *in vivo* demonstrate accumulation of senescence associated beta-galactosidase activity, DNA damage, and alterations to other senescence associated proteins^23,27^. In neurodegenerative diseases, some investigations have likewise shed light on the role senescent CNS cells have in mediating neurotoxic and injury related processes^33^. For instance, in Alzheimer’s disease (AD), the presence of senescent glia and neurons in the CNS are suspected to contribute to impaired memory and cell death, and clearance of senescent glia mitigates memory loss in AD transgenic mice^34,35^. Most recently, DNA damage driven gene fusions were detected in excitatory cortical neurons from AD patients and suggested to contribute to differential gene expression in neurons in the disease, including alterations to senescence pathways^36^. Similarly, glial and cortical neuron senescence has been reported following traumatic brain injury, where neurons accumulate DNA double strand breaks, loss of nuclear lamina integrity, and gene expression changes related to senescence processes^37-39^. Given the growing body of data that neuronal senescence occurs during aging and in response to various forms of injury and stress, it begs the question whether these characteristics of senescence arising in neurons is simply an outcome of accumulated injury over time, overwhelming the neuron’s intrinsic repair capacity, and impeding its homeostatic functions. Hence, further research into injury and stress driven senescence in neurons and how this may impede their normal function is still needed.

A fascinating piece of the puzzle is the role that transcription factors play in promoting neuronal resilience in response to injury and senescence. In our study, we found that overexpression of a combination of three transcription factors involved in cellular rejuvenation (*Oct4, Sox2, Klf4*) promoted neuronal survival in EAE. This approach was previously used to enhance survival and regeneration in RGCs in glaucoma and following acute injury in the optic nerve crush model through a *Tet1*-*Tet2* epigenetic modification dependent mechanism^20^. Thus, it seems rejuvenated neurons have potential for greater intrinsic injury response programs that enable them to survive, regenerate, and maintain function. Identifying drug combinations and therapies that can engage these programs is an exciting new approach as AAV-gene therapy, while a useful tool to study mechanisms, comes with numerous complications when applied in the clinic^40,41^. One possibility is the senomorphic drug metformin, which is currently being investigated for therapeutic potential in MS due to evidence that it may rejuvenate oligodendrocyte precursor cells and promote myelin repair^42,43^. It is conceivable that this approach may additionally promote a youthful profile in various cell types thereby exerting off target beneficial effects through cellular rejuvenation. Senolytics, a class of drugs designed to kill senescent cells, is another approach currently receiving attention for neurodegenerative diseases^44^. However, if neuronal senescence induces upregulation of the senolytic targets, these drugs may have the off-target and likely detrimental result of neuronal death. Importantly, treating senescence in MS requires careful consideration of the age of the patients as it is unique in the neurodegenerative disease field due to its typical onset in young adulthood. MS transitions from a predominantly neuroinflammatory condition in young people (20 to 30 years of age) to a progressive neurodegenerative disease in older patients, typically between 50 to 60 years of age^1,45^. Indeed, there is evidence of biomarkers of accelerated aging of the immune system, like shortened telomeres, that may contribute to the progressive decline in health experienced by older MS patients^45^. Thus, there remain many considerations and avenues for investigation into aging and senescence as a therapeutic target in the CNS in disease, particularly for MS. New approaches incorporating existing immunomodulatory agents, in combination with novel anti-senescence and promyelinating therapeutics may be key to promoting long term health of patients in the future^46,47^.

## Funding and acknowledgements

S.S.D., M.A.H., and A.M.R.G receive funding from the Canadian Institutes for Health Research (CIHR) Vanier Canada Graduate Scholarship. E.M.-L.H. is funded by the Fonds de Recherche du Quebec-Santé Master’s Training Scholarship. A.E.F. is funded by CIHR and MS Canada. D.A.S. is funded by NIH/NIA R01AG019719, a gift from Michael Chambers, and the Glenn Foundation for Medical Research to D.A.S who has equity and consults for Life Biosciences, a company developing rejuvenation medicines. For more information see https://sinclair.hms.harvard.edu/david-sinclairs-affiliations. We would like to acknowledge Dr. Stuart Trenholm for his expertise and support in the 3D printing and building of the optomotor response device. We would also like to acknowledge the essential technical support from Thomas Stroh and the Montreal Neurological Institute (MNI) Microscopy Core Facility, and Julien Sirois and the MNI Flow Cytometry Core Facility.

## Materials and methods

### Mouse husbandry

All animal experiments were performed according to the guidelines set by the Canadian Council on Animal Care (CCAC). Thy1-STP-YFP mice were bred with vGlut2-Cre mice to obtain mice expressing YFP in Thy1/vGlut2 expressing cells. These mice were kept and bred to maintain a Vglut2-Cre/Thy1-STP-YFP transgenic population.

### Experimental autoimmune encephalomyelitis (EAE)

EAE was induced 7–8-week-old female mice as described previously. Briefly, myelin oligodendrocyte protein (MOG) was prepared in an emulsion of complete Freund’s adjuvant (CFA) and 100 uL was injected subcutaneously. Two days later, animals were injected intraperitoneally with 400 ng pertussis toxin in 200 ul sterile Hank’s Balanced Salt Solution (HBSS). Animals were monitored for up to three weeks post induction for weight loss and motor symptom development. Motor symptoms were scored on a scale of 0-5, representing symptoms of ascending motor paralysis. All scoring was conducted blinded to treatment condition.

### Intravitreal injection of AAV virus

Bilateral intravitreal injections were performed three days prior to EAE induction. Briefly, mice were anesthetized with isoflurane, and the eye was treated with topical anesthetic for pain management during and after the procedure. Suture thread was used to pull back the eyelid and pull the conjunctiva to rotate the eye forward. A window was cut through the conjunctiva to visualize the injection location, then a microneedle was used to pierce the globe of the eye and inject the solution. The needle was held in place for 30 seconds to ensure mixing of the injected solution with the vitreous liquid, then the needle and suture thread were removed, and the mouse was monitored for recovery. 2 uL of virus mixture of AAV2-tTa-GFP/AAV2-CMV-tTa or AAV2-tTa-OSK/AAV2-CMV-tTa was injected per eye.

### Optomotor response

Optomotor response was tested in a custom-built drum with striped sheets rotated around the animal at 2 rpm in standard lighting^48^. Stripe sizes tested included 0.3, 0.35, and 0.4 cycles/degree (c/d) based on previous studies demonstrating a reduction in visual acuity from ∼0.4 c/d to 0.3 c/d during this timeframe of EAE. Animals were acclimated to the apparatus for 10 minutes per animal at least one day prior to collecting testing data. Animals that were unable to balance themselves on the platform were excluded from testing. Video data was analysed blinded to mouse identity and condition. Clockwise and counterclockwise responses were measured and compared as a group, such that each animal contributes two datapoints (one result per direction), since each eye is differentially affected in the disease course and contributes to the clockwise vs counterclockwise response.

### Tissue collection

Mice were perfused sequentially with ice-cold PBS and 4% paraformaldehyde (PFA). Eyes and optic nerves were dissected out and post-fixed in 4% PFA for 24 hours. The tissue was then rinsed with PBS and put in 30% sucrose for 24 to 72 hours, mounted in optimal cutting temperature reagent (OCT), flash-frozen with liquid nitrogen, and stored at -80 until cryo-sectioning. Retina sections were cut to a thickness of 12 um. For wholemount retinae and optic nerves, retinae were dissected out of the eye globe and both retinae and optic nerves were kept in PBS at 4°C until further processing.

### Retinal flat-mount immunohistochemistry

Retinae were washed three times with wash buffer (0.5% TritonX-100 in phosphate buffer solution (PBS)), then blocked and permeabilized overnight at 4°C in blocking buffer (5% normal donkey serum (DS), 2% TritonX-100 in PBS). Retinae were incubated in staining buffer (2% DS, 0.5% TritonX-100 in PBS) with primary antibodies for 72 hours at 4°C. Afterwards, retinae were washed 3 times for 10 minutes with wash buffer and incubated in secondary antibodies overnight in staining buffer. Retinae were washed again 3 times for 10 minutes with wash buffer, 1 time for ten minutes with PBS, then mounted on slides with Fluoromount G and left overnight at room temperature to cure.

### Retinal cross-section immunohistochemistry

Slides were thawed at room temperature for five minutes. For citrate antigen retrieval, slides were rehydrated first with ddH2O, then saturated with citrate buffer and heated to 90°C for ten minutes. Afterwards, slides were rinsed with room temperature citrate buffer. Slides were then washed three times for five minutes with 1X PBS, blocked and permeabilized in blocking buffer (5% DS, 0.2% TritonX-100 in PBS), and incubated overnight at 4°C in staining buffer (1% DS, 0.2% TritonX-100 in PBS) and primary antibodies. The next day, slides were washed three times for 5 minutes with 0.2% TritonX-100 in PBS, then incubated for 2 hours at room temperature in staining buffer with secondary antibodies. Slides were washed again three times five minutes with PBS, before mounting a coverslip, and curing overnight.

### Imaging and analysis

Images were taken on a confocal microscope with Zen software. Imaging parameters were kept the same between samples on the same slides. Automated analysis of nuclear fluorescence intensity was conducted for H3K27me3 in the RGC layer of cross-sections using a custom ImageJ macro. Counts of γH2AX+ RGCs and altered lamina RGCs were conducted manually, blinded to treatment condition. RBPMS+ RGC cell counts were conducted automatically using a custom ImageJ macro.

### Fluorescence activated cell sorting of RGCs

Mice were euthanized with CO2, then eyes were carefully removed with forceps. The retinae were immediately dissected out in cold Leibovitz-15 media and transferred to falcon tubes containing papain (10U/mL) and DNase in EBSS. Papain digestion was performed at 37°C for ten minutes, after which retinas were gently triturated to a single cell suspension. Ovomucoid solution was layered underneath the papain-single-cell suspension, and cells were spun down for 7 minutes at 1000 rpm. The papain/ovomucoid mixture was removed carefully as to not disturb the cell pellet, and then the pellet was resuspended in cold HBSS with 1% N2/B27 supplements and filtered through a 40-um mesh filter. Cells were stained with anti-CD90.2 and live-dead dye for thirty minutes, then spun down and resuspended in HBSS with 1% N2/B27 supplements. All data was acquired on the FACS Aria Fusion (Becton-Dicksinson Biosciences). Voltages were set up according to optimal PMT sensitive using the peak 2 (Spherotech) voltration technique. The nozzle used was 100 um at 20 psi. Doublet discrimination and dead cells were removed from sorting. Using the appropriate FMO control samples, a gate was drawn to select YFP/Thy1 double positive cells. Double positive cells were sorted into HBSS with 1% N2/B27 and held at 4°C until sort completion and downstream processing.

### ATAC-sequencing

Approximately 15 to 50 thousand cells were pelleted after FACS isolation by centrifuging at 500g for 5 minutes at 4°C. The pellet was carefully resuspended in 50 μL of cold lysis buffer (10 mM Tris-HCl, pH 7.4, 10 mM NaCl, 3 mM MgCl2, 0.1% IGEPAL CA-630), then immediately centrifuged at 4°C, 500g, for 10 minutes. The supernatant was pipetted off, and the pellet was resuspended in 50 uL of transposition buffer (25 uL Tagment DNA Buffer, 2.5 uL Tagment DNA enzyme 1, 22.5 uL nuclease free H2O) and incubated at 37°C for 30 minutes. After transposition, DNA was isolated using the Zymo ChIP DNA Clean & Concentrator Kit per manufacturers instructions, and eluted in 11 uL of 10 mM Tris Buffer, pH 8.0. Library preparation was done using Nextera primer sequences and NEB Next High Fidelity 2X PCR master mix (M0541L) and amplification was run for 10 cycles. 40 base-pair paired-end sequencing was performed on a NextSeq500, and sequences were demultiplexed with 0 mismatches using bcl2fast1 2.20.

### RNA-sequencing

Approximately 15 to 50 thousand cells were used per sample. Total RNA extraction was performed using the Qiagen miRNeasy kit according to manufacturer recommendations. 75 bp single-end sequencing was performed on a NextSeq500, and sequences were demultiplexed with 0 mismatches using bcl2fast1 2.20. Reads were trimmed with cutadapt, aligned with hisat2, and count matrices were generated by htseq. DESeq2 was used for count normalization and differential gene expression analysis. Pathway analysis was conducted with GSVA package for single-sample GSEA on the normalized counts table, and ClusterProfiler using the fgsea algorithm for ranked list GSEA^49,50^.

### Gene coexpression analysis

Gene co-expression analysis was conducted using SimpleTidyGeneCoEx workflow, which uses Leiden based clustering to group genes based on their common changes in expression across the sample sets^51^.

### Single-cell sequencing data analysis

The single-nucleus RNA sequencing (snRNA-seq) data in this study were obtained from a previously published dataset by Schirmer et al. in 2019, accessible under accession number PRJNA544731 (NCBI Bioproject ID: 544731)^52^. This dataset comprises gray matter tissue samples from eleven patients diagnosed with multiple sclerosis (MS) and nine control samples. The MS patient samples include those derived from acute-chronic active lesions and chronic inactive lesions of progressive forms of MS. To dissect the snRNA-seq data, we employed a well-established workflow with the Seurat package, encompassing processes such as dimensionality reduction, cell type identification, gene expression normalization, and clustering^53^. Specifically, cells with a mitochondrial gene composition of 5 percent or higher were classified as non-viable and were consequently excluded from further analysis. Likewise, cells with fewer than 200 or more than 2500 unique feature counts were deemed low-quality cells and were also removed from downstream analysis. Following quality control procedures, the gene expression profiles of the remaining cells were subjected to natural log normalization and scaling. Principal component analysis (PCA) was performed on highly variable genes to reduce the dimensionality of the dataset. Subsequently, the first 20 principal components were selected for clustering analysis. We constructed a shared-nearest neighbor graph based on the results of the PCA, and the Louvain clustering algorithm was employed iteratively to identify clusters at multiple resolutions. For this analysis, we used a resolution parameter of 0.6. Finally, we harnessed the uniform manifold approximation and projection (UMAP) algorithm to visualize the identified clusters in a two-dimensional space. Cell types were annotated using reference publication of Schirmer et al., datasets and clusters renamed based on their cell identities. To facilitate later analyses, we limited our analysis to neuronal cells with greater than 5 cells per cluster per condition (Control vs MS) in this dataset then we extracted and compiled the average expression levels of these neurons, stratified by patient.

